# The osteogenic potential of the neural crest lineage may contribute to craniosynostosis

**DOI:** 10.1101/321117

**Authors:** Daniel Doro, Annie Liu, Agamemnon E Grigoriadis, Karen J Liu

## Abstract

The craniofacial skeleton is formed from the neural crest and mesodermal lineages, both of which contribute mesenchymal precursors during formation of the skull bones. The large majority of cranial sutures also include a proportion of neural crest derived mesenchyme. While some studies have addressed the relative healing abilities of neural crest and mesodermal bone, relatively little attention has been paid to differences in intrinsic osteogenic potential. Here we use mouse models to compare neural crest osteoblasts (from frontal bones or dura mater) to mesodermal osteoblasts (from parietal bones). Using *in vitro* culture approaches we find that neural crest-derived osteoblasts readily generate bony nodules while mesodermal osteoblasts do so less efficiently. Furthermore, we find that co-culture of neural crest-derived osteoblasts with mesodermal osteoblasts is sufficient to nucleate ossification centres. All together, this suggests that the intrinsic osteogenic abilities of neural crest-derived mesenchyme may be a primary driver behind craniosynostosis.

---

Craniosynostosis (CS), the premature fusion of the cranial sutures, is a widely prevalent congenital malformation, affecting, on average, 1 in 2,000–2,500 live births^1^. The head in children with CS is often deformed, as a consequence of the compensatory growth at the remaining patent sutures, in an effort to accommodate the expanding brain^2^. In most cases, the anomalies are isolated (nonsyndromic); however, ∼25% of the defects are associated with over 180 different sydromes ^3,4^. In addition to the cranial and facial dysmorphisms that result from craniosynostosis, other morbities include increased intracranial pressure, potential developmental delay and, often, cognitive impairment^3^. Nonsyndromic craniosynostosis is multifactorial and most commonly affects the patency of sagittal or coronal sutures^5,6^. Metopic, lambdoid or multiple suture fusion accounts for less than 25% of nonsyndromic cranyosinostosis^7^.

Previous lineage tracing studies have described a predominance of neural crest derived mesenchyme in the large majority of the cranial sutures ^8–10^. Importantly, the maintenance of an undifferentiated and unossified mesenchyme is required for suture patency and often coincides with the establishment of a neural crest/mesoderm boundary, as clearly observed in the coronal suture, where frontal and parietal bones meet^11^. Moreover, abnormal proliferation and premature differentiation of the sutural mesenchyme are likely the most important cellular mechanisms leading to craniosynostosis^12^. These biological events are tightly regulated by intricate signalling pathways, originating from adjacent tissues with regional and temporal control over the suture fusion sequence^13^. For instance, the underlying dura mater has been implicated in specific determination of suture patency and ossification^14,15^

Rodents have been repeatedly used as a model for premature fusion of cranial sutures. Interestingly, only the posterior part of the frontal suture in mice fuses in early postnatal life, while all other cranial sutures remain patent throughout life^13^. Further studies in rats showed that a simple change of location to where the posterior frontal suture originally lies, was enough to induce the fusion of the sagittal sutures. The frontal suture, when transplanted to the anatomical position of the saggital suture, remained patent as the latter would^14^. The expression of osteogenic markers was potently upregulated in the dura mater underlying the closing posterior frontal suture, whereas the dura mater cells under the patent sutures show significantly increased proliferation^15^. The dura mater underlying the coronal suture was shown to be important for maintenance of patency, as removal of the membrane results in osseous obliteration in coronal suture transplants^16^. All together, these studies suggest that not only does the dura mater directly contributes osteoprogenitors to the ossification of the frontal suture, but the meninx juxtaposed to the cranial bones also provides regionally specified signals capable of inducing fusion of normally closing sutures, or maintance of patency in certain areas. The investigation of molecular events involved in physiological suture fusion is fundamental to the understanding of pathological premature obliteration as seen in craniosynostosis anomalies.

Dysfunctional suture fusion regulation performed by the dura mater has been previously proposed as an underlying cause for congenital craniosynostosis. Specifically, a colony of rabbits that models CS had the synostosed suture osteotomised and a patent suture transplanted to the site of the defect. The fusion of this suture was prevented by isolating the transplant from the underlying dura mater with an impermeable material, whereas synostosis occurred when no barrier was inserted ^17^. This suggests that the CS-inducing signal was coming from the underlying dura mater and not the from cranial bones or the sutures. Other studies have implicated fibroblast growth factor (FGF) as one amongst many potential dura mater-derived signals responsible for causing craniosynostosis^18^. Nonetheless, the origins and mechanisms behind craniosynostosis, as well as the osteogenic abilities of the distinct cranial populations still remain poorly understood.

Here we investigate the *in vitro* osteogenic capabilities of the distinct populations in the cranial complex, hypothesizing that the intrinsic potential of cells from different embryonic origins correlates with the potential for craniosynostosis. Ectopic suture fusion in ciliopathic mice, for instance, has been previously associated with aberrant expansion of the neural crest-derived tissues at the expense of mesodermal bones^19^. Utilising neural crest-specific lineage tracing techniques, we compared the bone forming ability of frontal and dura mater osteoprogenitors to mesodermally-derived parietal cells.

## Results

### Wnt1-Cre driver identifies the different embryonic origins of the cranial components

The embryonic origins of the mammalian skull have been previously mapped using the neural crest-specific *Wnt1-Cre* driver and the *Rosa26R^lacZ^* reporter^8,10^. To assess in detail the neural crest and mesodermal contributions to the cranial complex, we generated newborn pups (postnatal day 0, pN0) using the neural crest-specific *Wnt1-Cre* driver in association with a *Rosa26^mTmG^* reporter. In these mice, cells ubiquitously express a membrane Tomato (mT) marker. Upon expression of the Cre recombinase, cells switch from expression of membrane Tomato to membrane GFP (mG) specifically in neural crest tissues. We confirmed that most of the craniofacial skeleton expressed mGFP, indicating derivation from the neural crest lineage; one exception is the parietal bones, which continue to express membrane Tomato (mT) (Fig. 1a, b). As we dissected the cranial vault, we saw a predominant neural crest contribution to the sagittal suture, contrasting to a mixed contribution from neural crest and mesoderm to the coronal suture, where frontal and parietal bones meet (Fig. 1c, d). A closer look at the coronal suture reveals a clear neural crest-mesoderm boundary between frontal and parietal bones, which is underlined by a fully neural crest-derived dura mater. The overlying periosteum appears to be of mixed embryonic origin (Fig. 1e). Furthermore, we confirmed the distinct origins of the cranial components by enzymatically isolating cells from the postnatal bones and dura mater (PN0). After one day of culture, dura mater cells are more dense and elongated, whereas frontal bone cells appear more widely spread and sparse. Parietal bone cells resemble the morphology of frontal cells, although the well with the former was more densely populated and the cells smaller in size. (Fig. 1f, g, h). Although pure populations were rarely obtained when dissecting calvarial tissues, dura mater and frontal bone cells were largely green while parietal cells were red (Fig 1). Using this approach, these populations could be easily sorted according to their embryonic origins (Supplemental Figure 1). Because non-neural crest tissues (ectoderm, mesoderm and endoderm) could not be distinguished using the Wnt1-cre genetic model, we also confirmed the mesodermal origin of the parietal bones using the mesoderm-specific *Mesp1-Cre* driver (Supplemental Figure 2).

**Figure 1.**
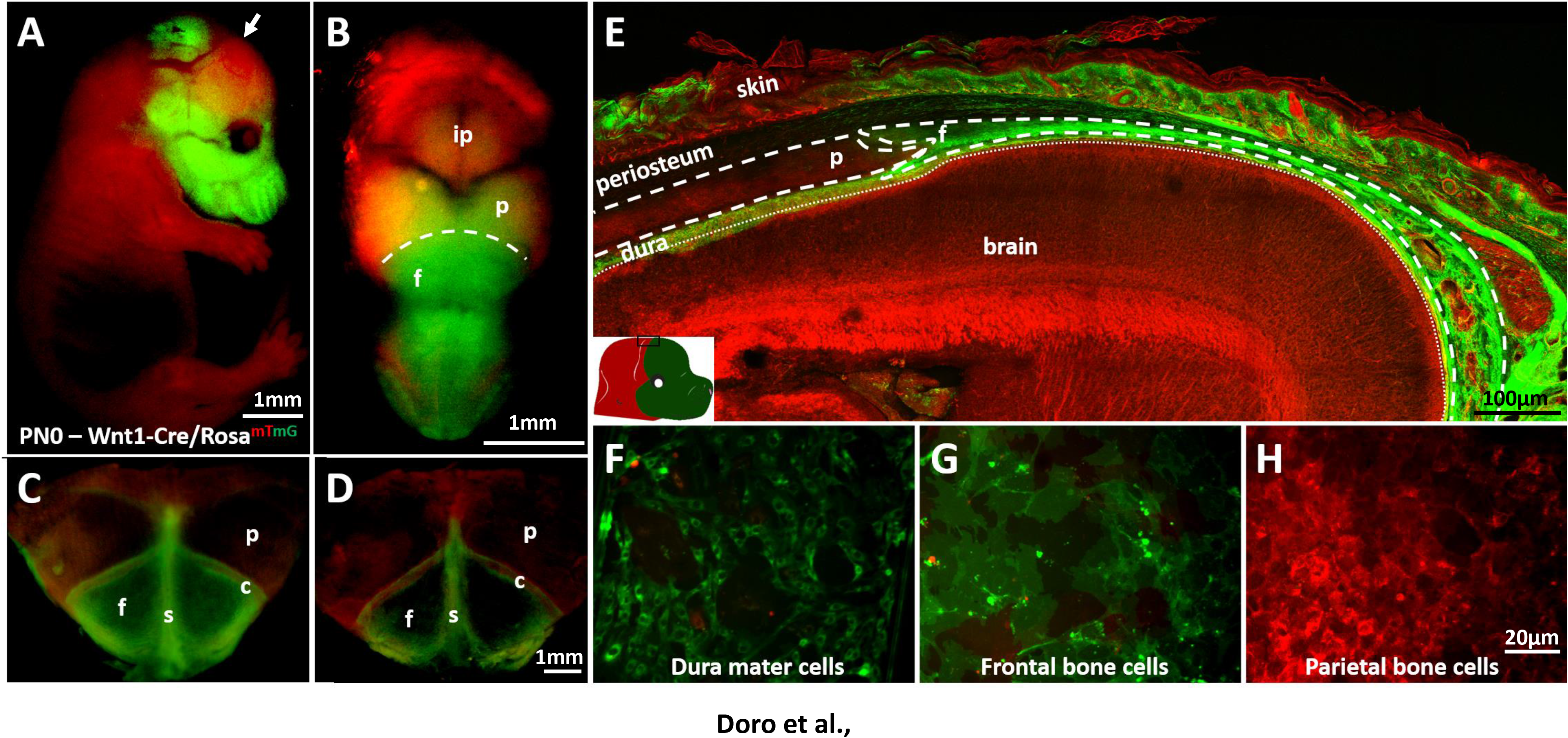
*Wnt1-cre*recombination labels neural crest-derived tissues in the cranial vault. (A-D) Whole mount views of new-born (PN0) *Wnt1-cre/Rosa^mT/mG^* tissues. (A) Side view of the whole animal reveals predominant neural crest origin (green) of the craniofacial structure with exception of the parietal region (arrow), which is mesodermally derived. All non-neural crest tissues appear in red. (B) Top view of the head shows neural crest-mesoderm boundary (dashed line) between frontal and parietal bones. Neural crest contribution (green) is also seen at the interparietal bone and on the midline between paired parietals. (C-D) Dissected cranial vaults with adjacent membranes intact (C), or after periosteum and dura mater removal (D) reveal the neural crest origin (green) of frontal bone and the whole sagittal suture and mesodermal contribution (red) of parietal bones and part of the coronal suture. (E) Sagittal section at the coronal suture shows the neural crest-mesoderm boundary in detail (boxed area in schematics). The neural crest frontal bone overlaps the mesodermally-derived parietal (dashed lines) at the coronal suture, whereas dura mater is exclusively neural crest. (F-G) Primary cells isolated from dissected calvarial tissues and cultured for 24 hours. (F) Dura mater and (G) frontal bone cells are neural crest derived (green) and (H) parietal cells are mesodermal in origin (red). (A-D) Scale bar = 1mm, (E) Scale bar = 100μm, (F-H) Scale bar = 20μm.f – frontal; p – parietal; ip – interparietal; c – coronal suture; s – sagittal suture.

**Figure 2.**
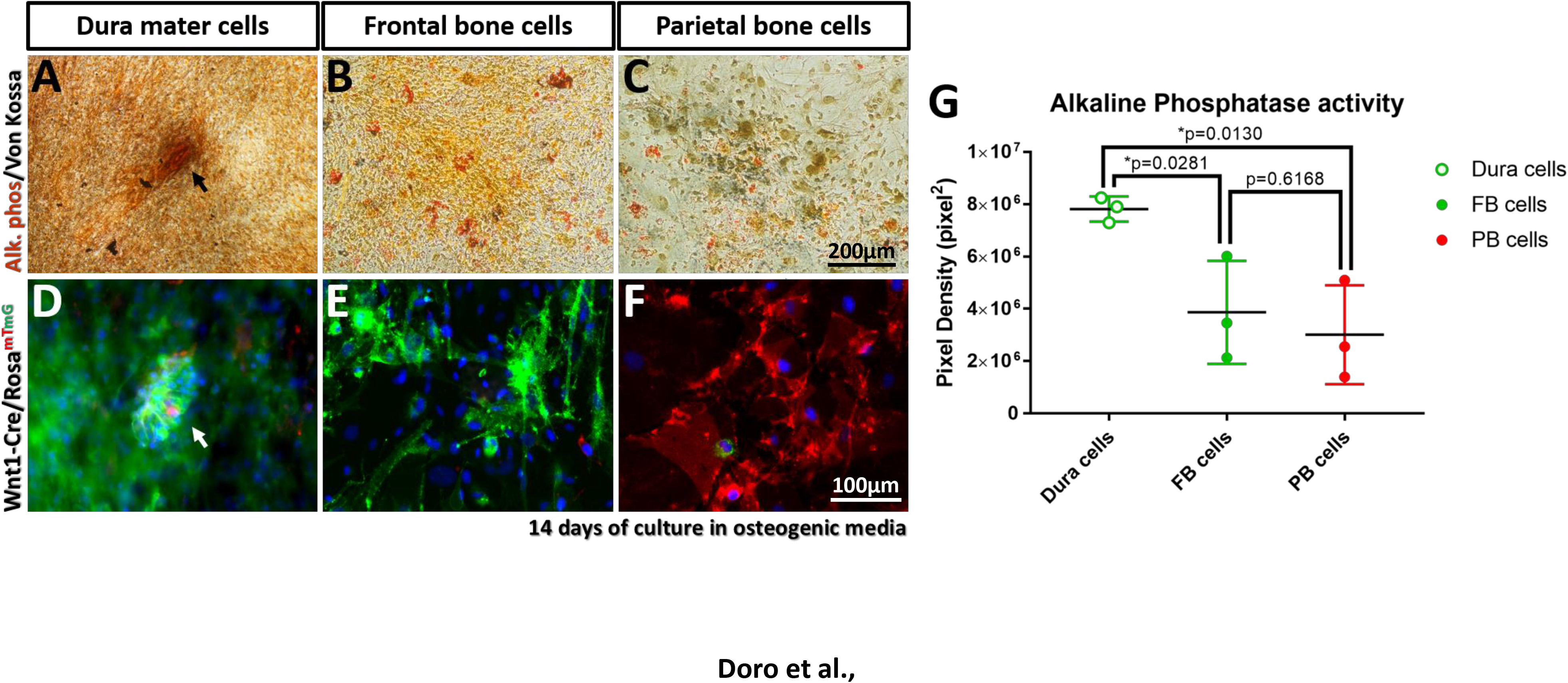
Dura mater cells are more osteogenic *in vitro* than osteoprogenitors from the cranial vault bones. (A-C) Alkaline phosphatase (ALP) (red) and Von Kossa (black) staining of PN0 cultured cells after 14 days in osteogenic media. (A) Dura mater cells were capable of making ALP positive, mineralized, bony nodules (arrow), whereas frontal (B) and parietal (C) cells are still sparse and ungrouped at this stage. (D-F) mGFP and mTomato fluorescent cells in higher magnification (Hoechst – nuclei). (D) Dura mater green cells are grouped in nodule-like structure (white arrow). (E) Frontal cells although sparsely distributed are more confluent than (F) parietal cells at this stage. (G) Alkaline phosphatase activity from calvarial cells, as measured by pixel density quantification of the red staining in A-C. Dura cells accordingly show significantly higher alkaline phosphatase activity when compared to frontal and parietal cells. (A-C) Scale bar = 200μm, (D-F) Scale bar = 100μm. \**p<0.05*. FB – frontal bone; PB – parietal bone.

### Neural crest-derived dura mater cells are more osteogenic in culture then frontal or parietal osteoprogenitors

To investigate the osteogenic potential of the isolated cell populations we cultured them at the same plating densities in osteogenic media. 14 days after plating, we observed the formation of mineralized nodules with intense alkaline phosphatase (ALP) activity in the dura mater cells (Fig. 2a and Supplemental Figure 3). Cells derived from frontal and parietal bones, on the other hand, do not make any nodules or aggregates at this stage (Fig. 2b, c). This is accompanied by significantly lower ALP activity, as assessed by the density of the staining (Fig. 2g). The fluorescence of the cells in culture revealed that while dura mater cells are in a stage of aggregation and nodule formation (Fig. 2d), frontal cells are still sparsely distributed and parietal cells are even more sparse and lower in density (Fig. 2e, f). This suggests that dura mater cells in culture have increased osteogenic potential in comparison to bone-derived osteoprogenitors. Moreover, a higher density and presence of aggregates in these cultures indicate an advanced stage of osteogenic differentiation, suggesting that frontal cells are at an intermediate stage between parietal cells and dura mater cells.

**Figure 3.**
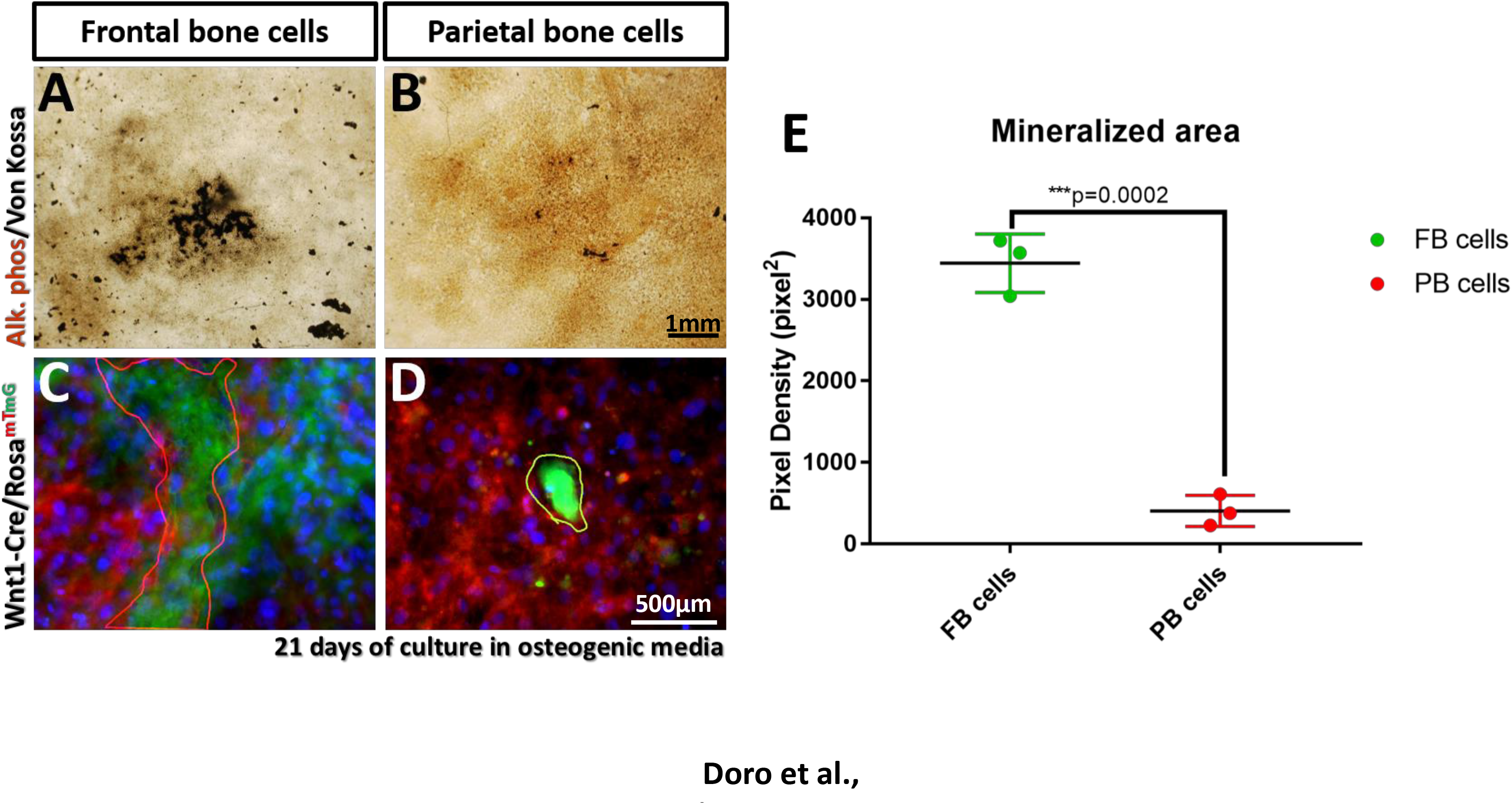
Neural crest frontal bone cells are more osteogenic *in vitro* than mesodermally derived parietal cells. (A-B) Alkaline phosphatase (ALP) (brown) and Von Kossa (black) staining of PN0 frontal and parietal cells after 21 days in osteogenic media. (A) While frontal cells were largely capable of making mineralized nodules, (B) cells isolated from the parietal bone show little mineralization and nodule formation. (C-D) mGFP and mTomato fluorescent cells in higher magnification (Hoechst – nuclei). (C) Even with the presence of non-neural crest contaminants (
red cells) in the frontal cells culture, the mineralized nodules (red outline) are mostly made of neural crest derived cells (green). (D) The rarely found nodules (yellow outline) in parietal cells culture seem to be made from neural-crest contaminants. (E) Mineralized area of frontal and parietal cell cultures, as measured by pixel density quantification of black staining in A-B. Frontal bone cells yield significantly more mineralized cultures than parietal cells. (A-B) Scale bar = 1mm, (C-D) Scale bar = 500μm. \*\*\**p<0.001*. FB – frontal bone; PB – parietal bone.

### Neural crest-derived frontal osteoprogenitors are more osteogenic than mesoderm-derived parietals in culture

To further confirm the increased potential of frontal bone cells over parietal cells, we cultured these populations at equivalent plating densities in osteogenic media for 21 days. This was enough to yield numerous mineralized frontal nodules, but very little mineralization of parietal cells (Fig. 3a, b). Indeed, the mineralized area was significantly higher in the former, as assessed by the density of (black) Von Kossa staining (Fig. 3e). Moreover, although the frontal bone cultures also contained cells of non-neural crest origin, nodules seen at 21 days were formed predominantly from neural crest derived progenitors (Fig. 3c). Interestingly, in the reciprocal experiment, the few nodule-like structures found in cultures of parietal cells appeared mostly green (Fig. 3d), suggesting a requirement of neural crest-derived progenitors for formation of the parietal bone condensations. This raises the hypothesis that parietal cells are generally incapable, or inefficient at making bony nodules *in vitro*. As these cells can ossify efficiently *in vivo*, it is likely that they are receiving pro-osteogenic signals from the adjacent neural crest.

### Co-culture with neural crest-derived cells favours in vitro osteogenesis of parietal osteoblasts

We next set out to test whether co-culturing of neural crest-derived cells and parietal progenitors at equivalent plating densities would produce bony nodules with mixed contributions. We observed that, after 14 days of culture, dura mater/parietal co-culture yields intense ALP activity and a large number of nodules (Fig. 4a). These nodules were mostly of mixed neural crest/mesoderm origin with no particular bias, although fluorescent labelling revealed that they appear to be nucleated by the green neural crest cells (Fig. 4c). Moreover, the culture of frontal bone cells with parietals resulted in vast mineralization and abundance of bony nodules after 21 days (Fig. 4b). Accordingly, these nodules were not skewed towards neural crest nor mesodermal cells (Supplemental Figure 4), although the base of the nodules was mostly formed by green cells (Fig. 4d). This was in contrast to the absence of mesodermally-derived nodules in culture of isolated parietal cells (Compare Fig. 4d and 3d). All together, our study indicates that, although mesodermally-derived parietal osteoprogenitors seem to lack the ability to efficiently make nodules in culture, they are able to contribute to the formation of these structures when co-cultured with cells of neural crest origin, which seem to nucleate the bony nodules. This raises the need to further investigate potential signals between calvarial cells of distinct embryonic origins, to improve our understanding of the molecular signalling at the neural crest-mesoderm boundary in the skull.

**Figure 4.**
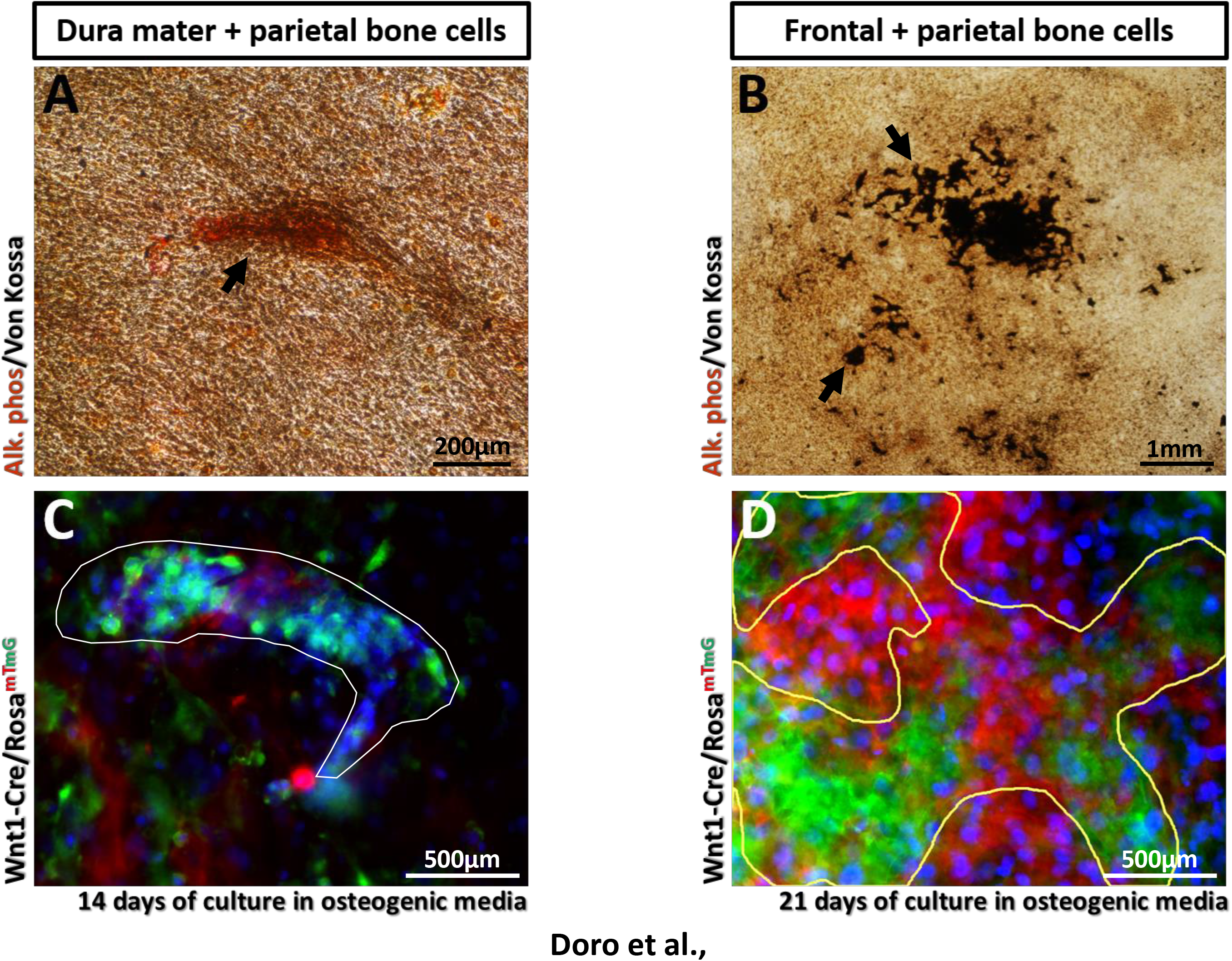
Parietal bone cells can contribute to nodule formation as long as they are co-cultured with neural crest derived cells. (A) ALP/Von Kossa staining of equivalently mixed dura mater and parietal cells reveal presence of mineralized nodules (arrow) after 14 days of culture in osteogenic media. (C) mGFP/mTomato fluorescence reveals mixed neural crest (green) and mesoderm (red) contribution for nodule (outline) formation (Hoechst – nuclei). (B) ALP/Von Kossa staining of frontal and parietal mixed cultures after 21 days in osteogenic media are largely mineralized (black staining). (D) mTmG fluorescence reveals neural crest (green) and mesoderm (red) mixed contribution for nodule formation (yellow outline). (A) Scale bar – 200μm, (B, D) Scale bar = 500μm, (B) Scale bar = 1mm

### Dura mater cells are the most osteogenic amongst neural crest derived cranial progenitors

In light of the higher propensity for osteogenesis displayed by the neural crest tissues in the cranial vault, we decided to investigate which of the site-specific progenitors would show higher osteogenic activity *in vitro*. PN10 osteoblasts were harvested from 3 distinct locations in Wnt1-Cre^mTdT/ +^ mice (see schematics on Figure 5L) and cultured for 21 days in osteogenic conditions. At day 21 we find that dura cells are denser and possess significantly higher alkaline phosphatase activity when compared to cells from the interfrontal suture (IFS) and frontal bone osteoblasts (FOBs) (Figure 5J). These cells also produce more mineralized nodules than the latter (Supplemental Figure 4b) and in larger size (Figure 5G-I). The presence of mineralized nodules is comparable between IFS cells and FOBs (Supplemental Figure 4G). However, the higher area of alkaline phosphatase staining indicates that frontal bone cultures have more progenitors committed to the osteogenic pathway, which are actively making bone *in vitro* (Figure 5A-C). This might reflect an expected delay in the differentiation stage, given that a proportion of the sutural mesenchyme is still uncommitted when harvested from the explants, whereas the cells harvested from the bone chips are likely fully differentiated at this postnatal stage. While it is true that the sutural mesenchyme harbours numerous undifferentiated stem cells with multi-lineage potential (reviewed in^20^), it is possible that, *in vitro*, these cells require exogenous signalling input to undergo osteogenesis. In addition, it would be interesting to know the dura mater requirements for ossification of the sutural cells.

**Figure 5.**
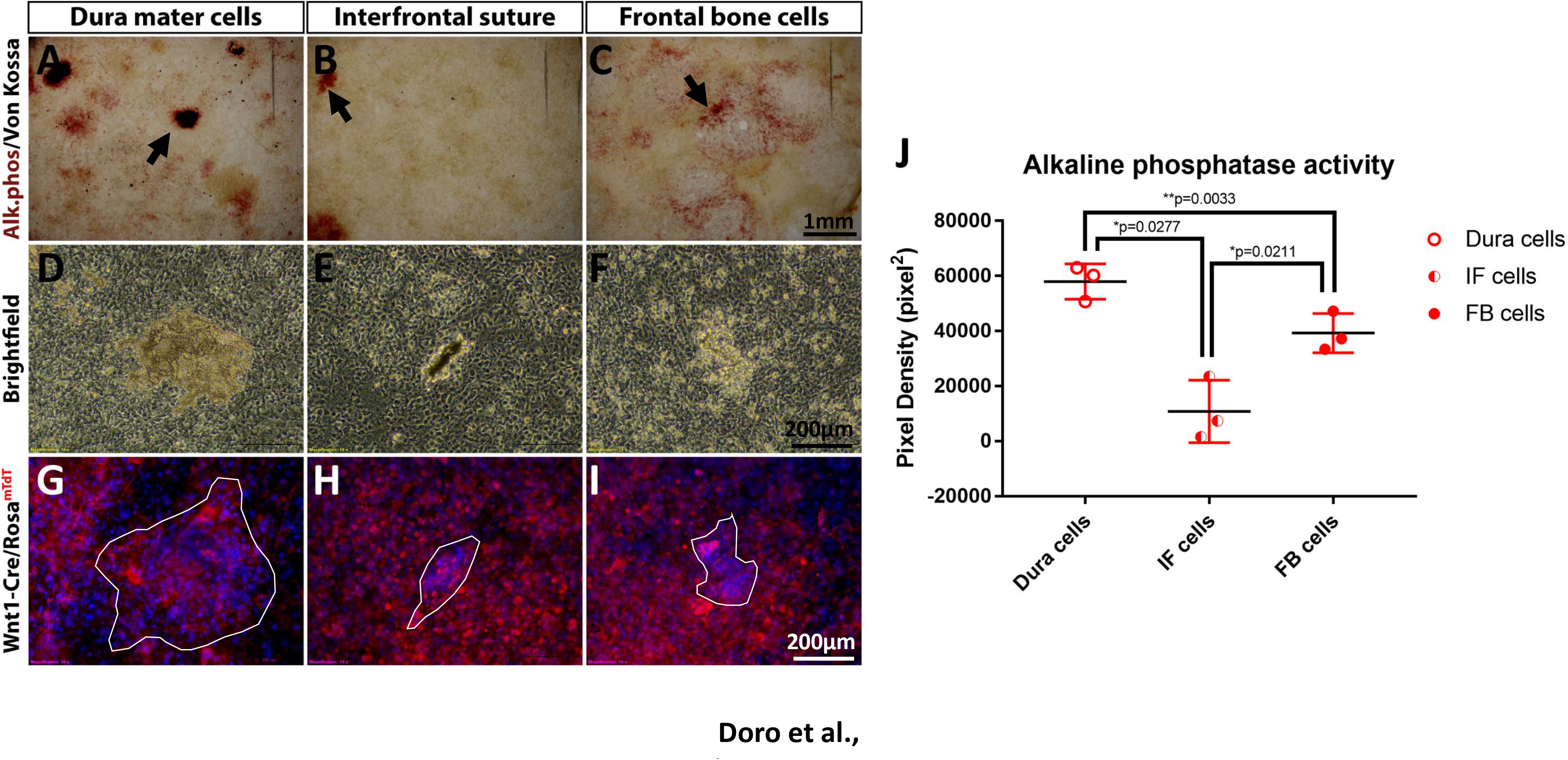
Dura mater cells are the most osteogenic amongst neural crest-derived cranial populations. (A-C) ALP/Von Kossa staining of primary cells from dura mater, interfrontal suture and frontal bone respectively. Arrows indicate mineralized nodules. (D-F) Contrast phase high magnification nodules reveal three-dimensional mineralized structure surrounded by a monolayer of cells. (G-I) mTomato fluorescence reveals neural crest origin of the Dura, IFS and FOB populations. The nodules are outlined and are increased in size in dura cell cultures. (J) Pixel density quantification of ALP staining shows significantly increased activity of dura cells when compared to IFS and FB cells, the latter is also significantly higher than in IFS cells. (L) Schematics show sites of harvesting where the cells originated from. (A-C) Scale bar = 1mm, (D-I) Scale bar = 200μm \**p<0.05, **p<0.005*. IF – interfrontal, FB – frontal bone.

## Discussion

The embryonic origins of the cranial components are often neglected in studies using primary cell cultures for osteogenic assays. While heterogenous populations isolated from whole calvaria have been largely used for *in vitro* differentiation assays, the bone forming ability of calvarial osteoprogenitors appears to be site- and lineage-specific. Each tissue in the cranial vault harbours distinct osteogenic capacity, being implicated in different roles during development and likely having distinct contributions to bone diseases and malformations. One classic example is the investigation of the pathogenesis of craniosynostosis.

Althought the premature ossification of a sutural mesenchyme may ultimately influence multiple tissues of distinct embryonic origins, it is still unclear which specific cell populations are the primary drivers of pathogenesis. Rather, as studies have shown, the dura mater seems to be the major player behind the suture patency, or fusion, during skull development. Either by providing molecular signals, or by directly contributing osteoprogenitors that undergo differentiation, the underlying dura mater may determine whether a cranial suture fuses or remains patent^15,21,22^. This membrane is regionally specified and can sometimes undergo aberrant differentiation, causing severe consequences for the adjacent tissue, as in craniosynostosis^17,18^, or even undergo tumorigenesis, as seen in meningeal osteosarcomas^23,24^. The understanding of the osteogenic potential and its correlation with the distinct embryonic origins is therefore fundamental for the comprehension of craniosynostosis and may reveal a propensity of neural crest-derived tissues to undergo hyperossification.

As we have reproduced in this study using genetic labelling and lineage tracing, the dura mater is exclusively neural crest-derived, as are the frontal bones. Even though the parietal bones are derived from the mesoderm, all the cranial sutures including the coronal are thought to be predominantly populated by neural crest osteoprogenitors. We have shown that the dura mater-derived cells are much more osteogenic *in vitro* than the osteoblasts harvested from frontal or parietal bones. This was revealed by an increased alkaline phosphatase activity and the prompt generation of bony nodules in dura mater cell cultures. Moreover, the osteoblasts derived from neural crest frontal bones showed significantly increased *in vitro* osteogenesis when compared to mesoderm-derived parietals. The latter were highly inefficient at making mineralized nodules in culture. However, the co-culture of parietal osteoblasts with neural crest-derived cells overcame this inefficiency, as the neural crest nucleated the bony nodules, allowing the mesodermal cells to engage in osteogenesis. This finding corroborates previous studies that report an ability of dura mater cells to provide a pro-osteogenic signal to cells in co-culture^17^.

The neural crest-derived cranial populations also showed distinct cell-autonomous osteogenic potential *in vitro*. We compared the alkaline phosphatase activity and the presence of mineralized nodules in cultures of frontal bone osteoblasts, dura mater and interfrontal suture progenitors. The dura mater, as expected, showed the highest osteogenic capacity, possibly accompanied by an increased proliferation, judging by the higher density of cells when compared to every other cranial population. Moreover, the cells derived from frontal bone osteoblasts displayed increased ALP activity in comparison to cells from the interfrontal suture. This could be indicative of a differentiation delay in the cells of the suture, which is unsurprising if we assume that the frontal osteoblasts are fully differentiated when harvested from the bone chips, while the sutural mesenchyme is still uncommited to the osteogenic lineage. Could the cells from the suture then be inherently programmed to remain undifferentiated in the absence of a pro-osteogenic cue? Do these cells require the input from the dura mater to undergo osteogenesis? Are any other lineages of a common progenitor obtained in these cultures at the expense of osteogenic cells? Future studies should include molecular analysis in order to confirm the differentiation status of the neural crest-derived cells.

The investigation of signalling cues is very important for understanding the pathophysiology of craniosynostosis. A recent study, in fact, has identified a number of *de novo* mutations in negative regulators of the Wnt, BMP and Ras/ERK pathways, occurring in non-syndromic midline craniosynostosis patients^25^. These pathways, knowingly implicated in positive regulation of osteogenesis, are, among other signals previously studied^18^, good candidates for dura mater’s regulatory milieu in the context of suture patency and fusion. All together, our study identifies distinct populations of osteoprogenitors in the cranial vault and provides a deeper understanding on the ossifying capabilities of these cells. Our hope is that this study will aid the comprehension of general suture biology and pathogenesis.

## Materials and Methods

### Animal Procedures

All animal work was performed at King’s College London in accordance with UK Home Office Project License P8D5E2773 (KJL). Mouse strains: Wnt1-Cre^26^, Mesp1-Cre^27^, R26R^mT/mG^ 28 or R26R^mTdT^ 29 mouse lines have all been described previously. Genotyping was performed as described in original publications. All the mouse lines have been bred on a mixed background, unless otherwise noted.

### Cryosectioning

Postnatal day 0 (PN0) mouse heads were harvested at birth. The samples were then fixed in 4% paraformaldehyde (PFA) for 48 hours at 4°C. After 3 1xPBS washing steps the heads were embedded in Optical Cutting Temperature (OCT) compound (CellPath®) in 3 steps. First, the samples were moved to a 30% sucrose solution in 1xPBS (∼24h). Then, the embedding solution was replaced with a 30% sucrose solution mixed with OCT compound (1:1) and incubated at 4°C. Finally, the samples were moved and oriented in a plastic Tissue-Tek^®^ Cryomold^®^ filled with OCT compound. The mold was quickly moved into a dry ice bath with absolute ethanol until the OCT block was fully solidified. OCT embedded samples were stored at –80°C and reconditioned to –26°C 30 minutes prior to sectioning. Cryosections were performed using OFT5000^®^ cryostat microtome (BrightInstruments®). 25μm sections were immediately mounted into Superfrost Ultra Plus^®^ slides (ThermoScientific®), which were then stored at room temperature for future use.

### Primary cell harvest

Dissections of postnatal calvaria were performed in 1x (PBS). Dura mater and periosteum were removed with a forceps prior to dissection of frontal and parietal. The sutures were trimmed off with micro scissors to avoid any mixing of the bone osteoblasts with mesenchymal progenitor populations. The mineralized matrix was then digested for 10 minutes with 0.5% Trypsin (Sigma-Aldrich), 20 minutes with 2mg/mL Dispase II (Roche) in Hank’s Balanced Salt Solution (HBSS) (Gibco) and 2 times 30 minutes with 2mg/mL Collagenase (Sigma-Aldrich) in HBSS. Each of the collagenase digestion steps was collected into tubes containing equivalent amount of Fetal Bovine Serum (FBS). After neutralization with FBS and mild centrifugation (1000rpm), the cell pellet was resuspended in 10mL of osteoblast culture media (see formula below) and plated into 10cm2 dishes. Plates were incubated at 37°C and 5% CO2.

Dura mater and interfrontal suture cells were obtained by plating pooled dissected dura mater membranes, or interfrontal sutures, after 10 minute digestion with 0.5% Trypsin (Sigma-Aldrich). The tissue was then incubated at at 37°C and 5% CO_2_ in osteoblast culture media.

### Fluorescence activated cell sorting (FACs)

Single calvariae, devoid of periosteum and dura mater, were digested according to the protocol above and the first and second collagenase pools were collected into tubes with FBS for neutralization. After centrifugation at 1000rpm, the pellet was resuspended in 500μL of PBS1x supplemented with 5% of FBS. The suspension obtained was pipetted through 40μm tip strainers (Flowmi™) to obtain a single cell suspension. The mixed mGFP and mTomato cell suspension was then sorted into green and red populations using BD FACS™Aria II sorter (BD^®^ biosciences).

### Cell culture

#### Osteoblast culture media

Minimum Essential Medium Eagle – Alpha modification (Alpha-MEM) with UltraGlutamineTMI, deoxyribonucleosides and ribonucleosides (Lonza®); 10% batch tested Fetal Bovine Serum and Gibco^®^ Antibiotic-Antimycotic (ABAM).

#### Osteogenic media

Alpha-MEM (Lonza®); 10% batch tested FBS; ABAM (Gibco®); 0.25mM ascorbic acid (Sigma®); 5mM β-glycerophosphate (Invitrogen®).

### Alkaline phosphatase staining

Alkaline phosphatase activity was detected after fixation of the cultured cells with 10% p-formaldehyde. After 3 1xPBS washes, the culture plates were incubated for 1 hour at room temperature in alkaline phosphatase staining solution: Sigma-Aldrich Fast Red Violet LB Salt (60mg) (Sigma®) and Naphtol AS-Mx-PO4 (10mg) (Sigma®) in 0.1M Tris pH 8.3 (100mL).

### Von Kossa staining

Von-Kossa staining was used for detection of mineralized nodules after fixation in para-formaldehyde and alkaline phosphatase staining. After 3 ddH_2_O washes, the culture plates were incubated in 2.5% of silver nitrate in water for 1 hour under UV exposure (252nm).

### Microscopy and Image Analysis

A stereoscope (Nikon SMZ1500) with an attached camera (Nikon digital sight DS-Fi1) was used to take all the whole mount pictures and fluorescent pictures at low magnification (as indicated). Confocal microscopy was performed on a Leica Microsystems CMS GmbH TCS SP5 DM16000. Image sequences were reconstructed using FIJI (Image J) analysis software. Pixel density analysis of staining was performed using the Analyze Particles function under the menu Analyse in the Image J software. This analysis is only possible after Adjusting the Threshold of the binary image created.

### Statistical analysis

P values were determined using multiple unpaired t-test analysis in the GraphPad Prism 7 Software. Every cell culture experiment was performed in triplicate, unless otherwise noted.

---

**Supplemental Figure 1.**
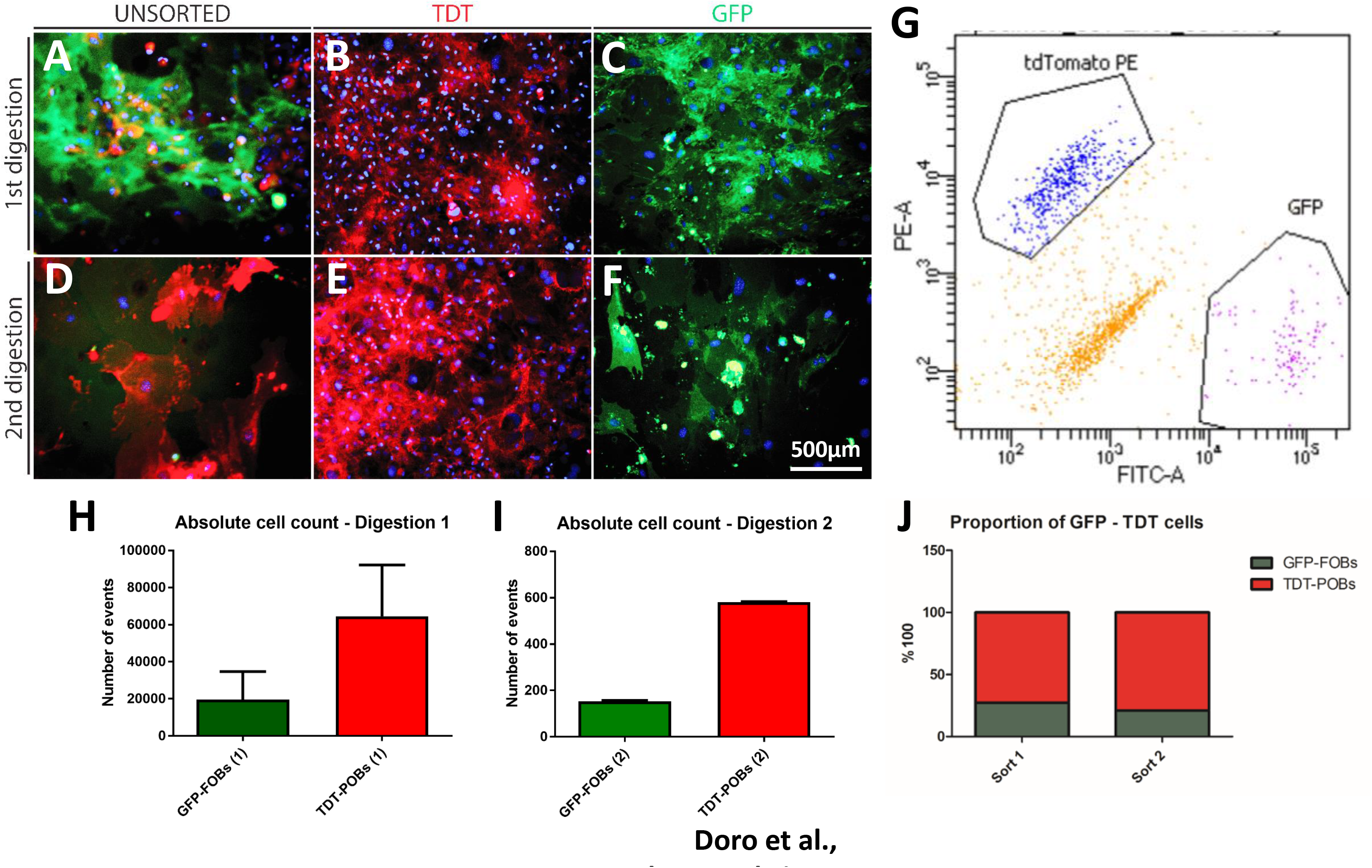
*Wnt1-cre/Rosa^mT/mG^* calvarial cells can be FAC sorted and suggest higher proportion of mesodermally derived parietal cells. (A-C) mTmG fluorescence of unsorted (A), red – mTomato (TDT) (B) and green – mGFP (C) calvarial cells digested in a first step from whole explants devoid of meninges and periosteum (see materials and methods). Blue – nuclear staining (Hoechst). (D-F) Cells from a second digestion are fewer in number and density. (G) Dot plot of Wnt1-Cre^mT/mG^ cells. mTomato positive cells (upper left) were sorted from mGFP positive cells (bottom right) and doubly labelled cells (lower left). Cells counted by the sorter reveal 3-fold higher proportion of red osteoblasts over green frontals in the first (H) and second (J) steps of enzymatic digestion. (A-F) Scale bar = 500μm. FOB – frontal osteoblasts, POB – parietal osteoblasts. n = 3 whole calvarial samples

**Supplemental Figure 2.**
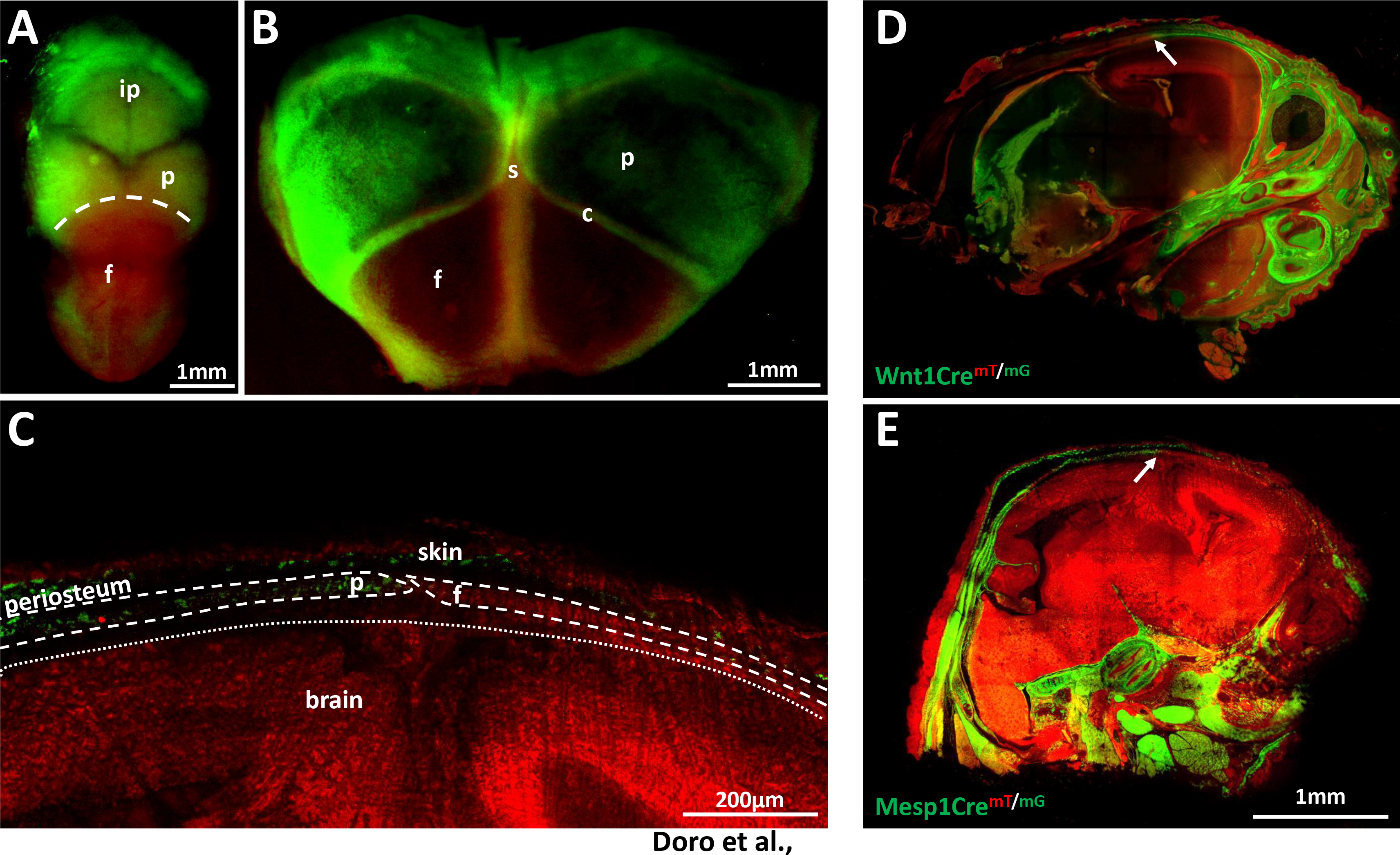
*Mesp1-cre* recombination labels mesoderm derived tissues in the cranial vault. (A) Top view of a PN0 Mesp1-Cre^mT/mG^ mouse reveals mesoderm derived tissues in green – mGFP. Parietal bones border with frontal bones (dashed line), which appear in red – mTomato as the rest of the face. (B) The dissected calvaria clearly shows the parietal bones originated from the mesoderm, which meet the frontal bones at the coronal suture region. (C) Sagittal section at the coronal suture shows the neural crest-mesoderm boundary in detail (bones outlined by dashed lines). The neural crest (red) frontal bone meets the mesodermally derived parietal (green) and is underlined by the neural crest dura mater (over dotted line). Panels D and E compare sagittal head sections of the neural crest and mesoderm specific lineage traced models Wnt1-Cre and Mesp1-Cre respectively. White arrows point to the coronal suture where the frontal bones meet the parietals. (A, B, D, E) Scale bar = 1mm, (C) Scale bar = 200μm. f – frontal, p – parietal, ip – interparietal, c – coronal suture, s – sagittal suture.

**Supplemental Figure 3.**
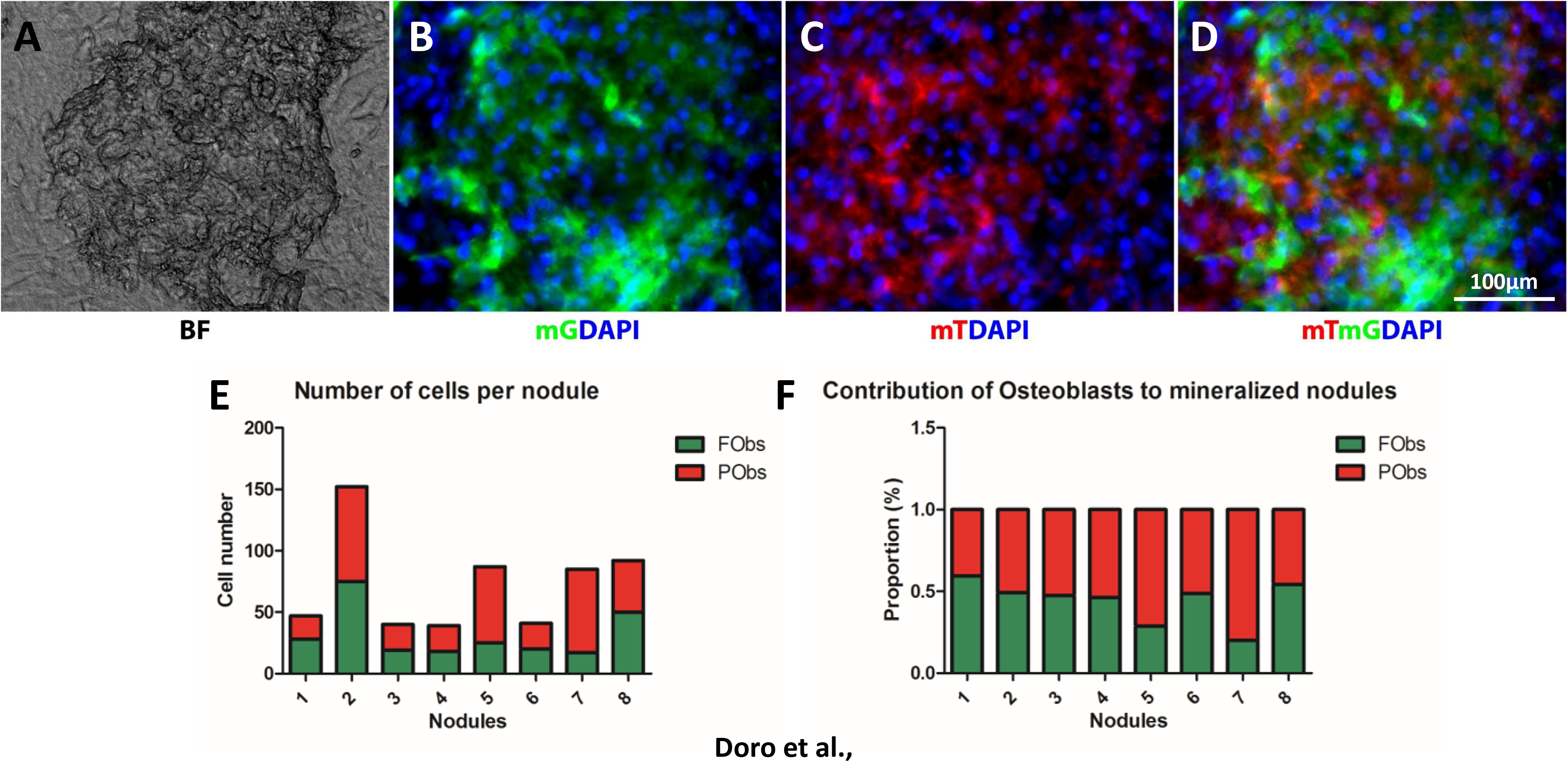
Co-culture of frontal and parietal osteoblasts generates nodules with equivalent proportion of neural crest and mesodermal cells. High magnification of a mineralized nodule from mixed populations of frontal – green and parietal – red osteoblasts. (A) Phase contrast shows three-dimensional structure composed of mGFP positive (B) and mTomato positive (C) cells. (D) Merged channels reveal that both cell populations contribute to the composition of the nodule. Blue – nuclei (Hoechst). (E) In terms of numbers, there is no particular prevalence of frontal (FObs) or parietal osteoblasts (PObs), as the bar chart shows. (F) Proportion in ratio (0.0–1.0) is variable and does not show more common prevalence of any specific population. (A-D) Scale bar = 100μm. mG – membrane GFP, mT – membrane Tomato.

**Supplemental Figure 4.**
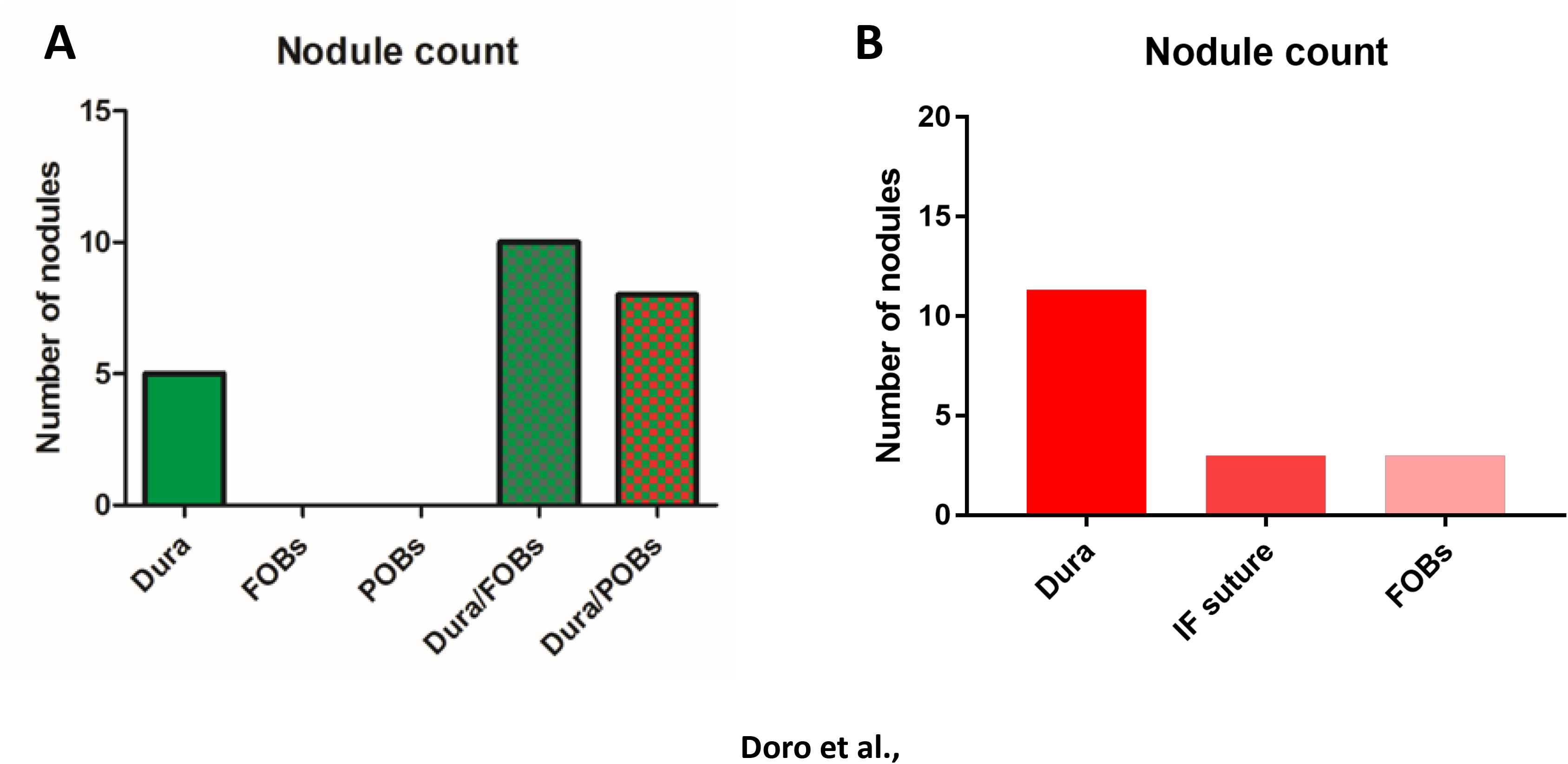
Dura mater cells produce more mineralized nodules after 2 weeks and 4 weeks of culture. Mineralized nodules were counted in lower magnification of each of 3 wells for every experiment. (A) Bar chart with average of nodules counted, after 14 day-incubation, in cultures of dura mater – green, frontal osteoblasts – dark green, parietal osteoblasts – red, 1:1 dura/FOBs co-cultured and 1:1 dura/POBs. While no nodule was seen in isolated cultures of frontal and parietal osteoblasts, upon co-culture with dura cells, these wells displayed a larger number of nodules than the dura cells cultured in isolation. (B) Average of nodules counted, after 21 day-incubation in cultures of isolated dura – bright red, interfrontal suture cells – dark pink and frontal osteoblasts – light pink. While the numbers of nodules seen in IF and FOBs are comparable, dura mater cultures yield a much larger number of mineralized nodules. FOBs – frontal osteoblasts, POBs – parietal osteoblasts, IF – interfrontal.

## References

1. Boulet SL, Rasmussen SA, Honein MA. A population-based study of craniosynostosis in metropolitan Atlanta, 1989–2003. Am J Med Genet Part A. 2008;146(8):984–991. doi:10.1002/ajmg.a.32208.

2. Morriss-Kay GM, Wilkie AOM.Growth of the normal skull vault and its alteration in craniosynostosis: insights from human genetics and experimental studies. J Anat. 2005;207(5):637–653. doi:10.1111/j.1469-7580.2005.00475.x.

3. Kimonis V, Gold JA, Hoffman TL, Panchal J, Boyadjiev SA. Genetics of Craniosynostosis. Semin Pediatr Neurol. 2007;14(3):150–161. doi:10.1016/j.spen.2007.08.008.

4. Greenwood J, Flodman P, Osann K, Boyadjiev SA, Kimonis V. Familial incidence and associated symptoms in a population of individuals with nonsyndromic craniosynostosis. Genet Med. 2014;16(4):302–310. doi:10.1038/gim.2013.134.

5. Hunter AGW, Rudd NL. Craniosynostosis. I. Sagittal synostosis; Its genetics and associated clinical findings in 214 patients who lacked involvement of the coronal suture (s). Teratology. 1976;14(2):185–193. doi:10.1002/tera.1420140209.

6. Hunter AGW, Rudd NL. Craniosynostosis. II. Coronal synostosis: Its familial characteristics and associated clinical findings in 109 patients lacking bilateral polysyndactyly or syndactyly. Teratology. 1977;15(3):301–309. doi:10.1002/tera.1420150312.

7. Lajeunie E, Le Merrer M, Marchac D, Renier D. Syndromal and nonsyndromal primary trigonocephaly: Analysis of a series of 237 patients. Am J Med Genet. 1998;75(2):211–215. doi:10.1002/(SICI)1096-8628(19980113)75:2<211::AIDAJMG19 > 3.0.CO;2-S.

8. Jiang X, Iseki S, Maxson RE, Sucov HM, Morriss-kay GM. Tissue Origins and Interactions in the Mammalian Skull Vault. 2002;116:106–116. doi:10.1006/dbio.2001.0487.

9. Yoshida T, Vivatbutsiri P, Morriss-Kay G, Saga Y, Iseki S. Cell lineage in mammalian craniofacial mesenchyme. Mech Dev. 2008;125(9–10):797–808. doi:10.1016/j.mod.2008.06.007.

10. Wu T, Chen G, Tian F, Liu H-X. Contribution of cranial neural crest cells to mouse skull development. Int J Dev Biol. 2017;61(8–9):495–503. doi:10.1387/ijdb.170051gc.

11. Merrill AE, Bochukova EG, Brugger SM, et al. Cell mixing at a neural crest-mesoderm boundary and deficient ephrin-Eph signaling in the pathogenesis of craniosynostosis. Hum Mol Genet. 2006;15(8):1319–1328. doi:10.1093/hmg/ddl052.

12. Iseki S, Wilkie AO, Morriss-Kay GM. Fgfr1 and Fgfr2 have distinct differentiation-and proliferation-related roles in the developing mouse skull vault. Development. 1999;126(24):5611–5620. http://www.ncbi.nlm.nih.gov/pubmed/10572038.

13. Bradley JP, Levine JP, Roth DA, McCarthy JG, Longaker MT. Studies in cranial suture biology: IV. Temporal sequence of posterior frontal cranial suture fusion in the mouse. Plast Reconstr Surg. 1996;98(6):1039–1045.

14. Levine JP, Bradley JP, Roth DA, McCarthy JG, Longaker MT. Studies in cranial suture biology: regional dura mater determines overlying suture biology. Plast Reconstr Surg. 1998;101(6):1441–1447. http://www.ncbi.nlm.nih.gov/pubmed/9583471. Accessed August 10, 2017.

15. Greenwald JA, Mehrara BJ, Spector JA, et al. Regional differentiation of cranial suture-associated dura mater in vivo and in vitro: Implications for suture fusion and patency. J Bone Miner Res. 2000;15(12):2413–2430. doi:10.1359/jbmr.2000.15.12.2413.

16. Opperman LA, Sweeney TM, Redmon J, Persing JA, Ogle ROYC. Tissue Interactions With Underlying Dura Mater Inhibit Osseous Obliteration of Developing Cranial Sutures. 1993;198312322.

17. Cooper GM, Durham EL, Cray JJ, et al. Tissue interactions between craniosynostotic dura mater and bone. J Craniofac Surg. 2012;23(3):919–924. doi:10.1097/SCS.0b013e31824e645f.

18. Ang BU, Spivak RM, Nah HD, Kirschner RE. Dura in the pathogenesis of syndromic craniosynostosis: Fibroblast growth factor receptor 2 mutations in dural cells promote osteogenic proliferation and differentiation of osteoblasts. J Craniofac Surg. 2010;21(2):462–467. doi:10.1097/SCS.0b013e3181cfe9a0.

19. Tabler JM, Rice CP, Liu KJ, Wallingford JB. A novel ciliopathic skull defect arising from excess neural crest. Dev Biol. 2016;417(1):4–10. doi:10.1016/j.ydbio.2016.07.001.

20. Doro DH, Grigoriadis AE, Liu KJ. Calvarial Suture-Derived Stem Cells and Their Contribution to Cranial Bone Repair. Front Physiol. 2017;8:956. doi:10.3389/fphys.2017.00956.

21. Warren SM, Greenwald JA, Nacamuli RP, et al. Regional dura mater differentially regulates osteoblast gene expression. J Craniofac Surg. 2003;14(3):363–370. http://www.ncbi.nlm.nih.gov/pubmed/12826808.

22. Mehrara BJ, Greenwald J, Chin GS, et al. Regional differentiation of rat cranial suture-derived dural cells is dependent on association with fusing and patent cranial sutures. Plast Reconstr Surg. 1999;104(4):1003–1013. http://www.ncbi.nlm.nih.gov/pubmed/106547401. Accessed April 14, 2018.

23. Bonilla F, Provencio M, Salas C, Espana P. Primary osteosarcoma of the meninges. Ann Oncol. 1994;5(10):965–966. doi:10.1093/oxfordjournals.annonc.a058744.

24. Ringenberg MA, Neitzel LE, Zachary JF. Meningeal osteosarcoma in a dog. Vet Pathol. 2000;37(6):653–655. doi:10.1354/vp.37-6-653.

25. Timberlake AT, Furey CG, Choi J, et al. De novo mutations in inhibitors of Wnt, BMP, and Ras/ERK signaling pathways in non-syndromic midline craniosynostosis. Proc Natl Acad Sci U S A. 2017;114(35):E7341–E7347. doi:10.1073/pnas.1709255114.

26. Danielian PS, Muccino D, Rowitch DH, Michael SK, McMahon AP. Modification of gene activity in mouse embryos in utero by a tamoxifen-inducible form of Cre recombinase. Curr Biol. 1998;8(24):1323–1326. doi:10.1016/S0960-9822(07)00562-3.

27. Saga Y, Miyagawa-tomita S, Takagi A, Kitajima S, Miyazaki J. MesP1 is expressed in the heart precursor cells and required for the formation of a single heart tube. 1999;3447:3437–3447.

28. Muzumdar MD, Tasic B, Miyamichi K, Li L, Luo L. A global double-fluorescent Cre reporter mouse. Genesis. 2007;45(9):593–605. doi:10.1002/dvg.

29. Madisen L, Zwingman TA, Sunkin SM, et al. A robust and high-throughput Cre reporting and characterization system for the whole mouse brain. Nat Neurosci. 2010;13(1):133–140. doi:10.1038/nn.2467.

